# Development of the pulmonary vasculature in the gray short-tailed opossum (*Monodelphis domestica*) – 3D reconstruction by microcomputed tomography

**DOI:** 10.1101/2024.04.19.590272

**Authors:** Kirsten Ferner

**Affiliations:** Museum für Naturkunde, Leibniz-Institut für Evolutions- und Biodiversitätsforschung, Invalidenstraße 43, 10115 Berlin, Germany

**Keywords:** marsupial, vascular genesis, µCT, lung, pulmonary artery, pulmonary vein

## Abstract

The marsupial gray short-tailed opossum (*Monodelphis domestica*) is born at the late canalicular stage of lung development and the lungs are structurally immature compared to eutherians. The majority of lung development, including the maturation of pulmonary vasculature, takes place in ventilated functioning state during the postnatal period.

The current study uses X-ray computed tomography (µCT) to three-dimensionally reconstruct the vascular trees of the pulmonary artery and pulmonary vein in the marsupial gray short-tailed opossum to investigate the vascular genesis during the postnatal period. The development of the pulmonary vasculature was examined in 15 animals from neonate to postnatal day 57. The final 3-D reconstructions of the pulmonary artery and pulmonary vein in the neonate and at 21, 35 and 57 dpn were transformed into a centerline model of the vascular trees. Based on the reconstructions, the generation of end-branching vessels, the median and maximum generation and the number of vessels were calculated for the lungs.

The pulmonary vasculature follows the lung anatomy with six pulmonary lobes indicated by the bronchial tree. The pulmonary arteries follow the bronchial tree closely, in contrast to the pulmonary veins, which run between the pulmonary segments. At birth the pulmonary vasculature has a simple branching pattern with a few vessel generations. Compared to the bronchial tree, the pulmonary vasculature appears to be more developed and extends to the large terminal air spaces. The pulmonary vasculature of the lungs from neonate to postnatal day 57 develops along the bronchial tree and shows a marked gain in volume and a progressive increase in vascular complexity and density. The gray short-tailed opossum resembles the assumed mammalian ancestor and is suitable to inform on the evolution of the mammalian lung. Vascular genesis in the marsupial bears resemblance to developmental patterns described in eutherians. Lung development in general seems to be highly conservative within mammalian evolution.

## Introduction

The mammalian lung needs to be adequately developed in utero to function as gas exchanging organ. Essential for a functioning lung are a very large surface area possessing a very thin air-blood barrier (Gehr et al., 1978), a mature surfactant system facilitating inflation stability (Clements, 1957; Orgeig et al., 2004; Sano and Kuroki, 2005), a tree-like system of conducting airways ventilating the gas exchange area (Storey & Staub, 1962; Tyler, 1983; Schittny, 2017) and an effective vascular system leading the blood to and from the gas exchange area (Hislop & Reid, 1972).

The formation and maturation of the blood vessels in the lungs is crucial for ensuring proper oxygenation of the blood and the overall function of the respiratory system. The lungs receive deoxygenated blood from the right ventricle via the pulmonary arteries and oxygenated blood from the left ventricle via the bronchial arteries. These pathways are usually termed pulmonary circulation/vasculature (vas publica) and bronchial circulation/vasculature (vasa privata), respectively (Mühlfeld et al., 2018). The present article will focus on the pulmonary vasculature.

The development of the pulmonary vasculature follows the course of mammalian lung development which can be categorized into five morphological stages (embryonic, pseudoglandular, canalicular, saccular and alveolar) (Copland & Post, 2004; Tschanz, 2007; Warburton et al., 2010; Schittny, 2017).

In the embryonic stage lung buds emerge from the foregut endoderm. The surrounding mesenchyme gives rise to the pulmonary vasculature (Lin et al., 2017). Cells from the cardiogenic mesoderm differentiate into cardiac progenitor cells, which give rise to the heart and the initial blood vessels. The cardiac tube is formed from the merging of the paired heart primordia. The heart undergoes partitioning into chambers, leading to the formation of the atria and ventricles.

In the pseudoglandular stage the preacinar branching pattern of airways and blood vessels is fully established (Jeffery, 1998). The Vascular Endothelial Growth Factor (VEGF) plays a crucial role in angiogenesis, promoting the growth of blood vessels in the developing lungs (Warburton et al., 2010). The pulmonary arteries and veins develop alongside the bronchial tree, ensuring that oxygen-depleted blood from the body is transported to the lungs for oxygenation and oxygen-rich blood is returned to the heart.

During the canalicular, saccular and alveolar stages, the respiratory tree is further expanded in diameter and length, accompanied by vascularization and angiogenesis along the airways. A massive increase in the number of capillaries occurs (Warburton et al., 2010).

After birth, changes in the circulatory system occur as the fetal shunts (ductus arteriosus and foramen ovale) close, redirecting blood flow to the lungs for oxygenation (Baudinette et al., 1988). The pulmonary vasculature continues to mature postnatally, with the formation of a dense capillary network around the alveoli to facilitate gas exchange (Schittny, 2017). Morphological studies on comparative lung development showed that the mammalian lung seems to be a highly conservative structure (Nakakuki, 1975, 1980; Szdzuy et al., 2008; Mess & Ferner, 2010; Ferner et al., 2017). The degree of lung maturation at birth and the timing of lung development might be different among mammalian species, but not the sequence of developmental steps resulting in final lung maturation.

Compared to eutherian neonates, which give birth to more developed offspring, marsupial neonates are born at a very early stage of development, often resembling embryos (Renfree, 2006). Essential organ systems of the marsupial neonate, including digestive, neuronal, immune and respiratory systems are immature and still under the process of development (Renfree, 2006; Szdzuy & Zeller, 2009; Ferner et al., 2017). Marsupial neonates are similar in development to a late eutherian fetus and the immature lung corresponds to the Carnegie stage 16-17 in the human fetus or E13-E14 in the fetal rat at (Wang et al., 2009; Fujii et al., 2020).

Several lung developmental studies in marsupial species have shown that the lungs of newborns are at the canalicular or saccular stage of lung development (Hill & Hill, 1955; Krause & Leeson, 1975; Gemmell & Little, 1982; Gemmell, 1986; Gemmell & Nelson, 1988; Runciman et al., 1996, 1998a; Frappell & Mortola 2000; Makanya et al., 2001, 2003, 2007; Burri et al., 2003; Szdzuy et al., 2008; Simpson et al. 2013; Modepalli et al., 2018; Ferner, 2021a). They are considered as functionally immature, resulting in a neonate with a limited respiratory performance that must rely to varying degrees on transcutaneous gas exchange (MacFarlane & Frappell, 2001; Simpson et al., 2011, 2013; Ferner, 2018, 2021b). The lungs of the newborn gray short-tailed opossum (*Monodelphis domestica)* are at the canalicular stage, characterized by a small number of large terminal air spaces providing insignificant surface area for respiration (Ferner, 2024).

Most of the studies of lung development in marsupial species have placed emphasis on the structural development of the gas exchange area (Krause & Leeson, 1975; Gemmell & Little, 1982; Gemmell, 1986; Gemmell & Nelson, 1988; Runciman et al., 1996, 1998a; Makanya et al., 2001, 2003, 2007; Burri et al., 2003; Szdzuy et al., 2008; Simpson et al. 2013), some provide information about airway structures (Krause & Leeson, 1973, 1975; Tucker, 1974, Cope et al., 2001; Cooke & Alley, 2002), but there is very little information about the developing pulmonary vasculature in marsupials (Baudinette et al., 1988; Bernard et al., 1993). Information about the pulmonary vasculature originate primarily from adult eutherians (Nakakuki, 1986, 1993, 1994 a, b; Kida, 1998; Counter et al., 2013; Mühlfeld et al., 2018). A few studies investigated the development of the pulmonary circulation, in particular of the pulmonary arteries, in rodent models (Razavi et al., 2012; Phillips et al., 2017) and in humans (Davies & Reid, 1970; Frey et al., 2004).

Traditionally the pulmonary vasculature of mammals was studied by producing celloidin or silicone rubber casts of the pulmonary artery and vein, mostly in combination with casts of the bronchial tree (e.g., Hislop & Reid, 1972; Nakakuki, 1986, 1993, 1994 a, b; Kida, 1998; Autifi et al., 2015). However, this technique is complicated by the pressure of the lung, which proves to be difficult to assess in small lungs, such as marsupials (Cooke & Alley, 2002). In recent years the development of new techniques, such as serial block-face scanning electron microscopy (SBF-SEM) (e.g., Mühlfeld et al., 2018; Buchacker et al., 2019) or µCT in combination with 3-D remodeling (e.g., Vasilescu, 2012; Counter et al., 2013; Razavi et al., 2012; Ackermann et al., 2014; Phillips et al., 2017) opened new possibilities to examine and describe the three-dimensional structure of the mammalian pulmonary vasculature and alveolar capillary network. However, a three-dimensional reconstruction of the pulmonary vasculature in a marsupial species is missing so far. The only study examining the developing lung of two marsupial species by phase contrast imaging methods resulted in three-dimensional volume rendering of the air spaces, but neglected the bronchial tree and pulmonary vasculature of the lung (Simpson et al., 2013).

The gray short-tailed opossum offers a unique opportunity for a better understanding of the development of the mammalian pulmonary vasculature given the unique finding that, in contrast to eutherian mammals, the entire process of postnatal lung development including the formation of the pulmonary vasculature occurs in a ventilated functioning state. Since the gray short-tailed opossum resembles both the supposed marsupial and mammalian ancestor (Szdzuy & Zeller, 2009; Deakin et al., 2013; Ferner et al., 2017), it is suitable to inform on the evolution of the mammalian lung.

As a first step, the development of the bronchial tree and of the terminal air spaces of the lung of *Monodelphis domestica* were investigated (Ferner and Mahlow, 2023; Ferner 2024). The present study was targeted at the development of the pulmonary vasculature of the lung of the gray short-tailed opossum during the postnatal period using microcomputed tomography (µCT). In addition, we aimed to obtain functional volumes of the pulmonary vasculature using three-dimensional (3D) reconstructions of computed tomography data.

## 2. Material and methods

### 2.1. Specimen collection

The gray short-tailed opossums (*Monodelphis domestica*) used for this study were obtained from a laboratory colony established at the Museum für Naturkunde Berlin. After controlled mating, the females were checked for offspring when approaching full-term (13-14 days). 15 animals ranging from neonate (defined as the first 24 h at the day of birth) to 57 postnatal days were collected, weighed and euthanized by anesthetic overdose with isoflurane under animal ethics permit approved by the Animal Experimentation Ethics Committee (registration number: T0202/18). The specifics and numbers of the specimens are summarized in Table 1.

**Table 1.** Gray short-tailed opossum (*Monodelphis domestica*) specimens examined in this study. Body weights, lung volumes and volumes of pulmonary artery and pulmonary vein are presented.

| Age (days) | Species number | Body weight (g) | medium | Staining | Lung volume (mm<sup>3</sup>) | Bronchial tree volume (mm<sup>3</sup>) | Pulmonary artery volume (mm<sup>3</sup>) | Pulmonary vein volume (mm<sup>3</sup>) |
| --- | --- | --- | --- | --- | --- | --- | --- | --- |
| neonate | 1965_1* | 0.12 | Araldite | - | 2.70 | 0.21 | 0.06 | 0.06 |
|  | 2350_1 | 0.13 | Karnovsky | PTA | 1.73 | 0.09 | 0.08 | 0.08 |
|  | 2350_7 | 0.13 | Karnovsky | PTA | 2.74 | 0.24 | 0.08 | 0.03 |
|  | mean (SD) | 0.13 (0.01) |  |  | 2.39 (0.57) | 0.18 (0.08) | 0.07 (0.01) | 0.06 (0.03) |
| 4 dpn | 1995_5 | 0.22 | Bouin/70% Eth. | Iodid | 4.10 | 0.16 | 0.10 | 0.06 |
|  | 2257_6 | 0.21 | Karnovsky | PTA | 4.19 | 0.23 | 0.09 | 0.11 |
|  | 2257_8 | 0.21 | Karnovsky | PTA | 5.60 | 0.24 | 0.12 | 0.10 |
|  | mean (SD) | 0.21 (0.01) |  |  | 4.63 (0.84) | 0.21 (0.04) | 0.10 (0.02) | 0.09 (0.03) |
| 7dpn | 1987_5 | 0.34 | Bouin/70% Eth. | PTA | 6.08 | 0.42 | 0.11 | 0.07 |
|  | 2383_2 | 0.28 | Karnovsky | PTA | 5.63 | 0.68 | 0.14 | 0.08 |
|  | mean (SD) | 0.31 (0.04) |  |  | 5.86 (0.32) | 0.55 (0.18) | 0.13 (0.02) | 0.08 (0.01) |
| 11dpn | 1993_2 | 0.69 | Bouin/70% Eth. | Iodid | 20.58 | 1.00 | 0.30 | 0.18 |
| 14dpn | 1994_9 | 0.98 | Bouin/70% Eth. | PTA | 24.87 | 2.10 | 0.45 | 0.42 |
| 21dpn | 2040 | 2.43 | Bouin/70% Eth. | PTA | 71.14 | 4.00 | 1.86 | 1.54 |
| 28dpn | 2059 | 4.22 | Bouin/70% Eth. | PTA | 189.75 | 13.08 | 4.68 | 3.15 |
| 35dpn | 2065 | 7.25 | Bouin/70% Eth. | Iodid | 388.59 | 28.71 | 8.97 | 9.03 |
| 49dpn | 2049 | 13.64 | Bouin/70% Eth. | Iodid | 504.03 | 25.38 | 14.97 | 13.65 |
| 57dpn | 2179 | 31.58 | Bouin/70% Eth. | Iodid | 1176.68 | 103.51 | 28.20 | 29.01 |

dpn, days post natum; Eth., Ethanol; PTA, phosphotungstic acid; \* Specimen processed for Electron microscopy

### 2.2. Lung fixation

Young ranging from neonate to 28 days post natum (dpn) were decapitated, to allow better lung fixation via the trachea, and the whole animals were immersed in Karnovsky fixative (2g paraformaldehyde, 25ml distilled water, 10 ml 25% glutaraldehyde, 15 ml 0.2 M phosphate buffer) or Bouin’s solution (picric acid, formalin, 100% acetic acid, 15 : 5 : 1; Mulisch & Welsch, 2015). The fixation time in Bouin was usually one to two days. Afterwards the specimens were rinsed in 70 % ethanol. Specimens fixed with Karnovskys fixative stayed in the fixative until scanning (between some weeks and months). In late developmental stages, from 35 dpn to 57 dpn, the lungs were fixed via the trachea. The animal was placed in a supine position and the trachea was exposed via a midline incision, a cannula with polyethylene catheter tubing of an appropriate outer diameter was introduced into the trachea and the lungs were then fixed via the trachea with Karnovsky solution at a pressure head of 20 cm, until fixative was emerging from nostrils and mouth. Cardiovascular perfusion was not performed to preserve the original structural organization and to avoid artificial dilatation of the blood vessels. The small size and delicateness of the lung and cardiac system, especially in the early developmental stages, made it impossible to perfuse the cardiovascular system. In small developmental stages (0 - 28 dpn) the entire torsos were scanned in µCT. In older developmental stages the lungs and hearts were dissected together for µCT-scanning.

One neonate was fixed for transmission electron microscopy (TEM). The specimen was cut in two halves and the upper part of the trunk was processed for TEM. For this purpose, the specimen was fixed in 2.5% glutaraldehyde buffered in 0.2 M cacodylate (pH 7.4) for 2 hours, rinsed with 0.1 M cacodylate buffer, postfixed in 1% osmium tetroxide and embedded in epoxy resin (Araldite).

### 2.3. Preparation for µCT

µCT-imaging of soft-tissue structures has been limited by the low intrinsic x-ray contrast of non-mineralized tissues. For visualization of the lungs and pulmonary vasculature in µCT the specimens had to be stained in advance to impart sufficient contrast to these soft tissues.

With very simple contrast staining, µCT imaging produces quantitative, high-resolution, high-contrast volume images of lung tissue. This is possible without destroying the specimens and offers the possibility to combine it with other preparation and imaging methods (histology or TEM).

Metscher (2009) summarized several simple and versatile staining methods for µCT-imaging of animal soft tissues. Based on this information, different staining protocols using inorganic iodine and phosphotungstic acid (PTA), were developed, tested and used to produce high-contrast x-ray images of the lung and pulmonary vasculature at different age stages (Table 1). Staining with tungstic acid (PTA) was either performed in ethanol with 1% PTA for 21 to 42 days (full body specimens of 14 to 28 dpn) or in an aqueous solution (Karnovsky fixative) starting with 0.5% for 7 to 20 days and increased afterwards to 1% resulting in a staining period up to 30 days (Metscher, 2009). Separated lungs of older stages (35-57 dpn) were stained in 1% Iodine (Gignac et al., 2016) to test staining differences and shrinking effects, which could not be detected. The differences in staining periods and staining concentration depended on the respective specimen size and preparation. Torsos and lungs were scanned in distilled water using a small container. The specimens were fixed in the container with cotton balls to avoid moving around during the scan.

The specimen processed for electron microscopy could be scanned without further processing. The postfixation with osmium tetroxide leads to well-contrasted scanning results.

### 2.4. µCT imaging

The prepared specimens were subjected to micro-tomographic analysis at the Museum für Naturkunde Berlin (lab reference ID SCR_022585) using a Phoenix nanotom X-ray|s tube (Waygate Technologies, Baker Hughes, Wunstorf, Germany; equipment reference ID SCR_022582) at 70–110kV and 75–240μA, generating 1440–2000 projections (Average 3– 6) with 750–1000ms per scan or a YXLON FF85 (equipment reference ID SCR_020917) with transmission beam for bigger specimens at 90–110kV and 100–150μA, generating 2000 projections (Average 3) with 250-500ms. The different kV, µA and projection-settings depended on the respective machine and specimen size, which is also responsible for the range of the effective voxel size between 1.5–20.1μm. The cone beam reconstruction was performed using the datos|x 2 reconstruction software (Waygate Technologies, Baker Hughes, Wunstorf, Germany; datos|x 2.2).

### 2.5. Segmentation, visualization and data analysis for 3D reconstruction

The 3D volume processing was done with the software Volume Graphics Studio Max Version 3.5 (Volume Graphics GmbH, Heidelberg, Germany). µCT data were analyzed in detail as serial two-dimensional (2D) and reconstructed to three-dimensional (3D) images (Fig. 1). 3 D reconstructions of the bronchial tree and of the terminal airspaces were carried out before (Ferner & Mahlow, 2023; Ferner, 2024).

**Fig. 1.**
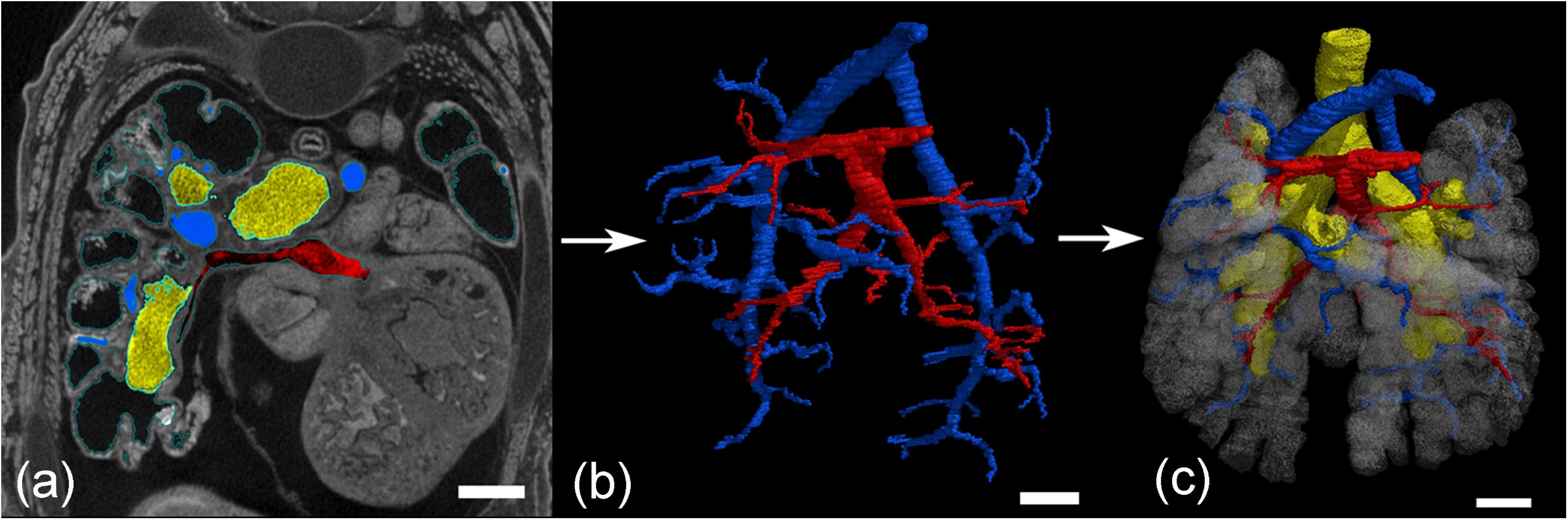
Three-dimensional reconstruction of the bronchial tree using 3D X-ray µCT. (a) Representative 3D X-ray µCT image in the transverse section in a neonate (2350_7). (b) The pulmonary artery (blue) and pulmonary vein (red) were reconstructed. (c) Pulmonary vasculature together with bronchial tree (yellow) and terminal airspaces (transparent white). Scale bar = 0.3 mm.

The segmentation was performed on 16-bit images to reconstruct the entire pulmonary artery and pulmonary vein respectively. A region grower tool was used, that marks all areas of the same density-value connected to each other to create a region of interest (ROI). The tissue density is mapped to gray values, so that tissues of the same density appear in the same gray scale value. A tolerance of 1000-1200 gray scale values around the first selected gray value of the ROI (center of the pulmonary artery or vein) was given. The reconstruction of the pulmonary artery (marked in blue colour) started from the centerline of the pulmonary trunk, coming from the right ventricle. The reconstruction of the pulmonary veins (marked in red colour) started at the point where the pulmonary veins enter the left atrium of the heart. The region grower tool was extended to the vessel walls. From there the ROIs were extended by scrolling through the image stack and applying region growing to the walls of the pulmonary artery and veins respectively until the blood vessels / capillaries became too small for reconstruction. It was visually ensured that only blood vessels were included. In that way the bronchial tree, the terminal air spaces and other air-filled areas in or between the lung segments were excluded from the segmentation.

The calculations of ROI-volumes are built-in functions of Volume Graphics Studio Max. The volumes of the ROIs of the pulmonary artery and pulmonary vein were calculated by the program Volume Graphics and indicated by mm³. The values are presented in Table 1. The volumes of the entire lung, the bronchial tree and the terminal air spaces were published before (Ferner and Mahlow, 2023; Ferner 2024). The final 3 D reconstructions of the pulmonary artery and pulmonary vein in the neonate and at 21, 35 and 57 dpn were transformed into a centerline model of the vascular trees visually showing the generation of the branches (Fig. 8). Based on the centerline model, the median and maximum generation and the number of vessels were calculated for the left, right and total lung and the values are presented as single values or mean and standard deviation (Table 2).

**Table 2.** Development of the branching tree of the pulmonary artery and pulmonary vein in *Monodelphis domestica* from neonate to 57 dpn.

| Age (days) | Median of Generation |  |  |  |  |  | Maximum Generation |  |  |  |  |  | Number of vessels |  |  |  |  |  |
| --- | --- | --- | --- | --- | --- | --- | --- | --- | --- | --- | --- | --- | --- | --- | --- | --- | --- | --- |
|  | Right lung |  | Left lung |  | Total lung |  | Right lung |  | Left lung |  | Total lung |  | Right lung |  | Left lung |  | Total lung |  |
|  | PA | PV | PA | PV | PA | PV | PA | PV | PA | PV | PA | PV | PA | PV | PA | PV | PA | PV |
| Neonate 1 | 7 | 4.5 | 4 | 4 | 5 | 4 | 10 | 8 | 6 | 6 | 10 | 6 | 24 | 18 | 10 | 7 | 41 | 17 |
| 21 dpn 1 | 6 | 6 | 6 | 7 | 6 | 6 | 10 | 9 | 9 | 12 | 10 | 12 | 34 | 24 | 29 | 41 | 56 | 70 |
| 35 dpn 1 | 10 | 9 | 7 | 8 | 10 | 8 | 20 | 16 | 11 | 14 | 20 | 14 | 141 | 61 | 70 | 48 | 202 | 118 |
| 57 dpn 1 | 16 | 12 | 12 | 11 | 14 | 12 | 27 | 24 | 25 | 25 | 27 | 25 | 404 | 292 | 358 | 286 | 695 | 644 |

dpn, days post natum; PA, pulmonary artery; PV, pulmonary vein

## Results

### General anatomy of the pulmonary vasculature

The lung of *Monodelphis domestica* has six pulmonary lobes. The right lung consists of separate cranial, middle, caudal and accessory lobes and the left lung consists of fused middle and caudal lobes. The pulmonary vasculature follows the lung anatomy indicated by the bronchial tree closely.

The right and left pulmonary arteries arise from the pulmonary trunk, which originates from the right ventricle of the heart. The pulmonary trunk bifurcates into right and left pulmonary arteries below the arch of aorta. The right and left pulmonary arteries cross the ventral surface of their respective main bronchus and curve laterally then dorsally to attain a position dorsal to each bronchus. The right and left pulmonary arteries enter the lung at the hilum along with the main bronchi. After entering the lungs, the pulmonary arteries run in parallel to the bronchial tree. As the right and left main bronchi divide into lobar bronchioles, the right and left pulmonary arteries divide into lobar branches that follow the bronchial division. In the left lung two major lobar branches of the pulmonary artery communicate with the middle and caudal lobes. The middle lobar branch originates ventrolateral from the left main pulmonary artery and divides into two arteries which supply the cranial and caudal parts of the left middle lobe. The large second branch of the left pulmonary artery enters the caudal lobe as the caudal lobar branch and sends branches along the caudal lobar bronchial division. In the right lung, the main pulmonary artery divides into cranial, middle, accessory and caudal lobar branches, corresponding to the pulmonary lobes. The cranial lobar branch originates from the dorsolateral surface of the right main pulmonary artery. The cranial lobar bronchiole arises above the level of the right pulmonary artery, and for this reason is named the eparterial bronchiole. The middle lobar branch originates from the ventrolateral surface of the right main pulmonary artery. The accessory lobar branch originates ventromedially from the right pulmonary artery. From this point the pulmonary artery continues caudally as caudal lobar branch which extends with many branches along the caudal lobar bronchial division of the right lung. In general, the lobar arteries course along the dorsal surface of the lobar bronchioles. Only the accessory lobar artery passes between the middle and caudal lobar bronchioles of the right lung to course along the ventral surface of the accessory lobar bronchiole.

The pulmonary veins can be distinguished in three main pulmonary veins, two emerging from the hilum of the right lung (cranial lobe vein and a main pulmonary vein (middle pulmonary vein)) and one main pulmonary vein coming from the left lung. The right, middle and left pulmonary veins, which return oxygenated blood from the lungs to the left atrium of the heart, are formed by the pulmonary lobar veins which receive blood from several feeding veins coming from each pulmonary lobe. The pulmonary lobar veins lay along the ventral surface of the corresponding lobar bronchiole except for the accessory pulmonary lobar vein which is dorsal to the accessory lobar bronchus.

The right cranial pulmonary vein lies in front of and below the pulmonary artery. The right pulmonary vein is formed by the right cranial pulmonary lobar vein and the right middle pulmonary lobar vein. In addition, an extra vein from the cranial part of the right caudal lobe may contribute to the right pulmonary vein. The right caudal pulmonary lobar vein and the accessory pulmonary lobar vein join to form a middle pulmonary vein. The left pulmonary vein is formed by a branch from the cranial part of the cranial lobe and a branch from the caudal part of the cranial lobe which join the left caudal pulmonary lobar vein. The right, left and middle pulmonary veins join to form a common pulmonary venous trunk that opens into the left atrium of the heart.

Generally, the pulmonary veins are running on the ventromedial side of the main bronchi, distantly following the bronchial tree. In contrast to the pulmonary arteries, the peripheral feeding veins do not follow the bronchial tree closely. They run between the pulmonary segments from which they drain the blood.

Representative reconstructions of the pulmonary vasculature of all developmental stages of *Monodelphis domestica* are shown in Figure 2 and 3. The course of the pulmonary vasculature in relation to the bronchial tree viewed from different perspectives is presented in Figure 4, 5 and 6. Details of the developing pulmonary vasculature are given in Figure 7. The generations of end-branching vessels for a neonate, and animals aged 21, 35 and 57 dpn are presented in Figure 8 and the median, maximum and total number of vessel generations are summarized in Table 2. The volumes of the lung, the bronchial tree, the pulmonary artery and pulmonary vein are plotted against body mass (Fig. 9a) and the volumes of the pulmonary artery and pulmonary vein are plotted against postnatal age (Fig. 9b).

**Fig. 2.**
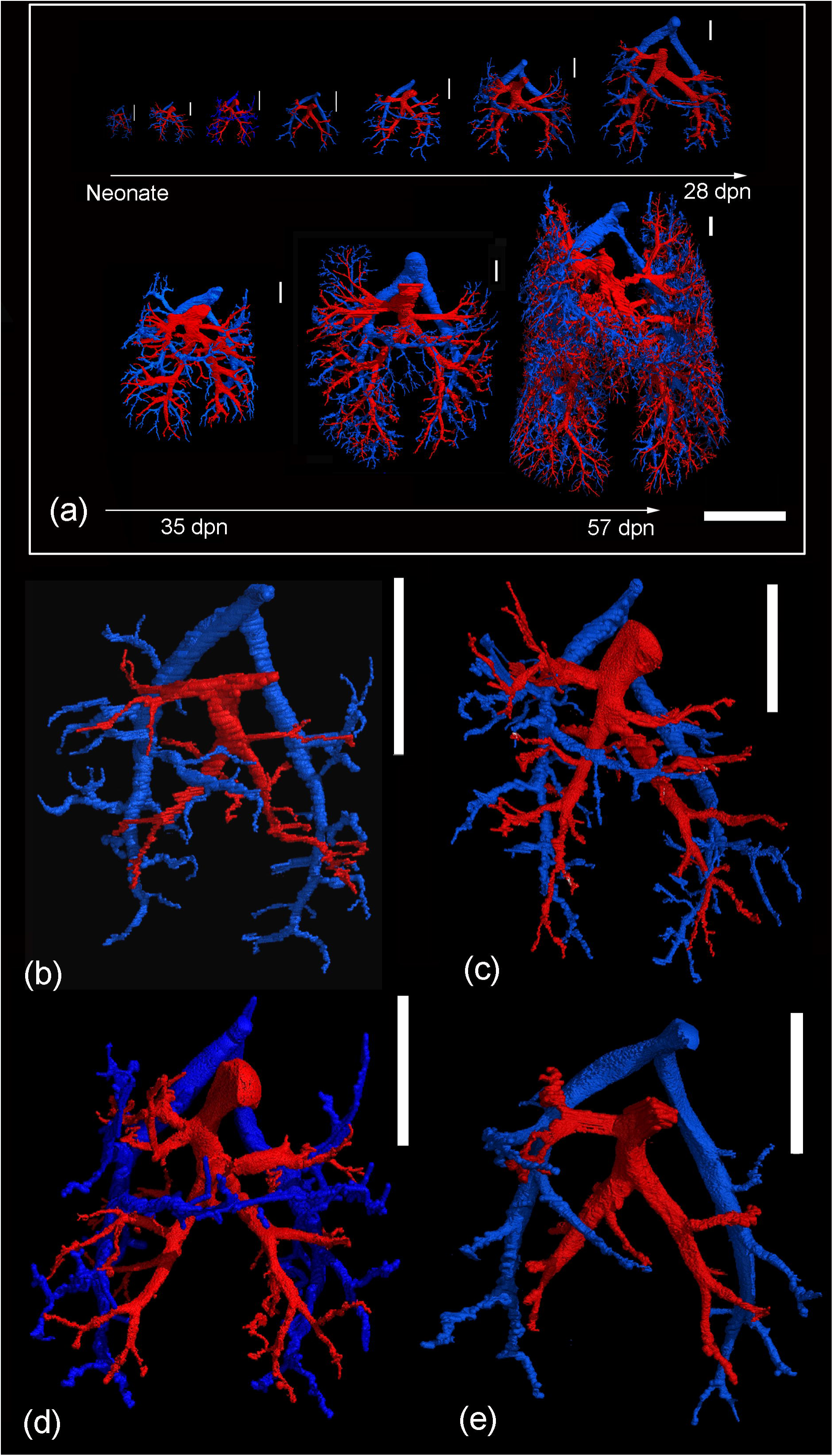
3D reconstructions of the pulmonary vasculature according to growth presented with the same scale highlight the impressive size increase of the developing lung from neonate to adult (a). Representative reconstructions of the pulmonary vasculature of *Monodelphis domestica* in ventral view in the neonate (b), at 4 dpn (c), at 7 dpn (d) and at 11 dpn (e). The pulmonary artery and vein are indicated by blue and red respectively. The scale bar is 5 mm in (a) and 1 mm in (b-e).

**Fig. 3.**
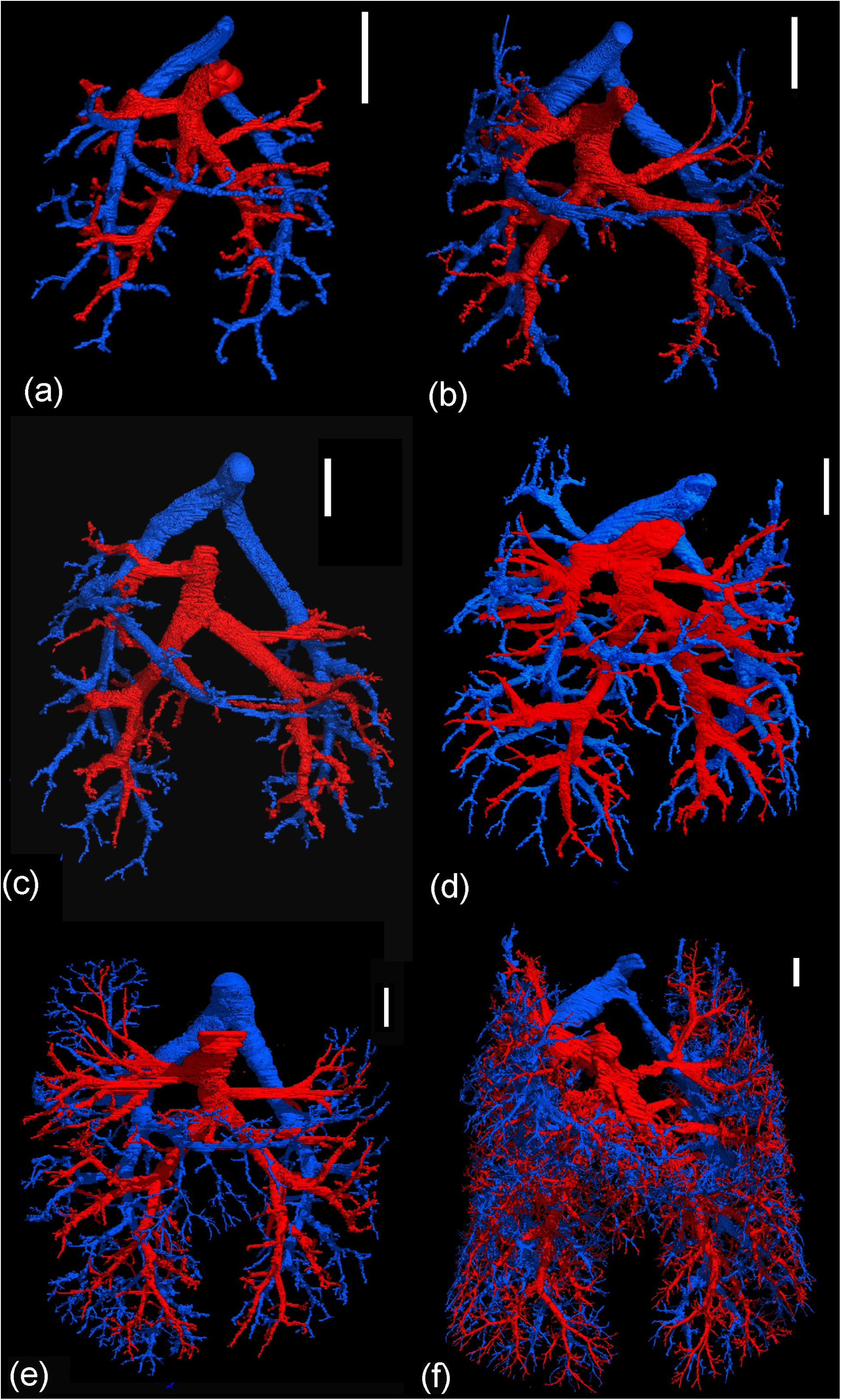
Representative reconstructions of the pulmonary vasculature of *Monodelphis domestica* in ventral view at 14 dpn (a), at 21 dpn (b), at 28 dpn (c), at 35 dpn (d), 49 dpn (e) and at 57 dpn (f). The pulmonary artery and vein are indicated by blue and red respectively. The scale bar is 1 mm.

**Fig. 4.**
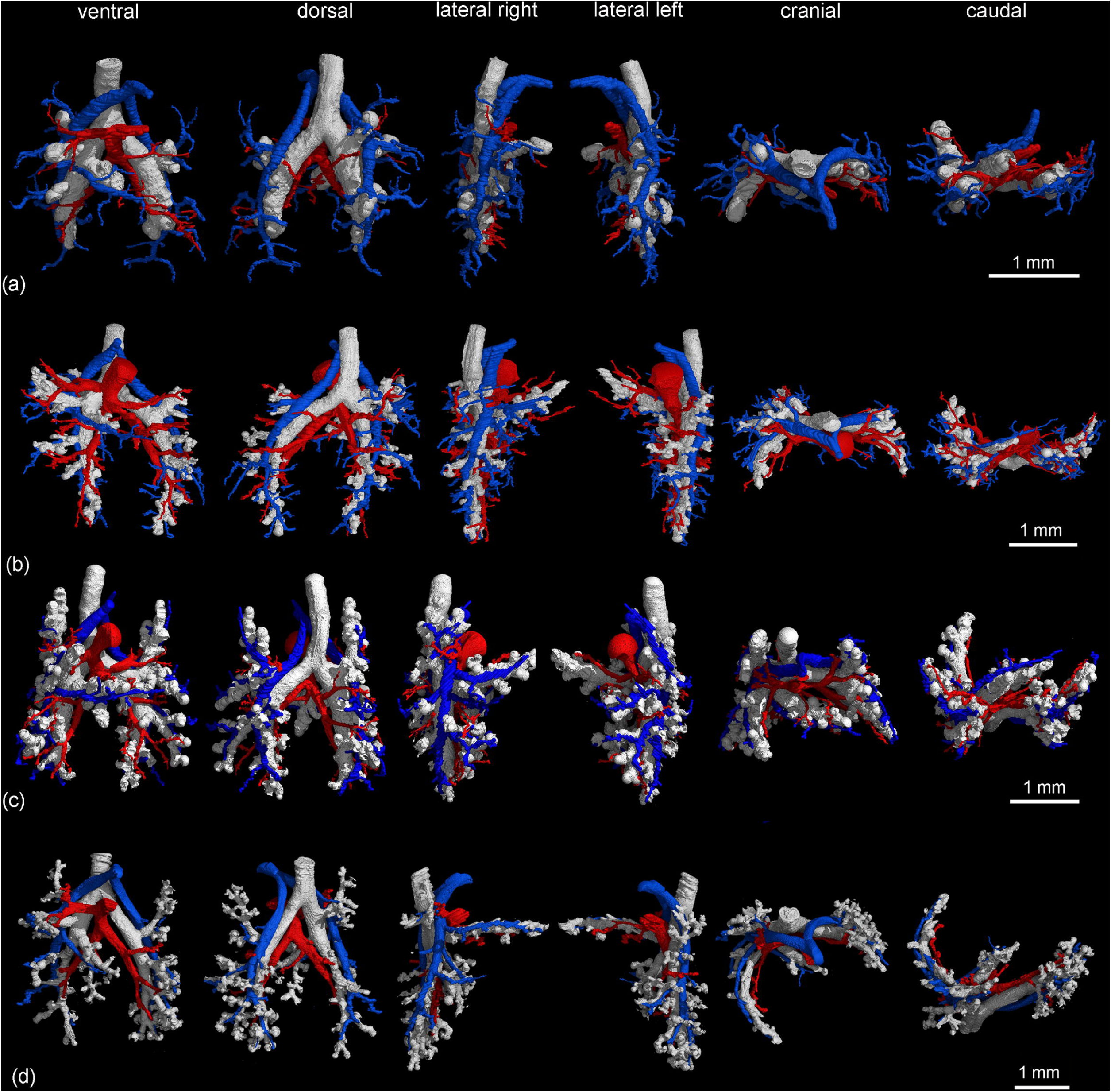
Pulmonary vasculature in relation to the bronchial tree in the neonate (a), at 4 dpn (b), at 7 dpn (c) and at 11 dpn (d). The lungs are shown from different perspectives: in ventral, dorsal, lateral, cranial and caudal views (from left to right). The pulmonary artery and vein are indicated by blue and red respectively, the bronchial tree appears white.

**Fig. 5.**
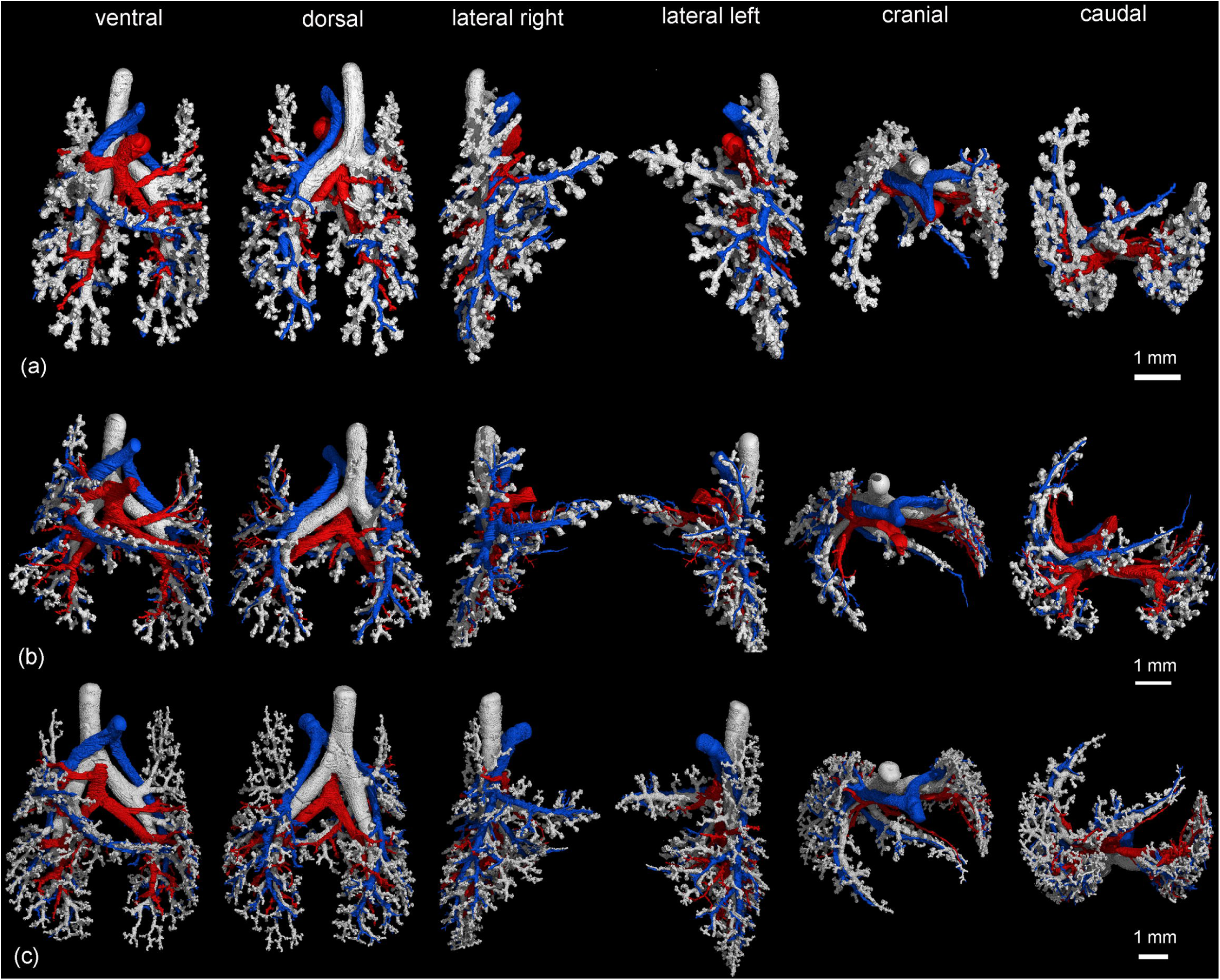
Pulmonary vasculature in relation to the bronchial tree at 14 dpn (a), at 21 dpn (b) and at 28 dpn (c). The lungs are shown from different perspectives: in ventral, dorsal, lateral, cranial and caudal views (from left to right). The pulmonary artery and vein are indicated by blue and red respectively, the bronchial tree appears white.

**Fig. 6.**
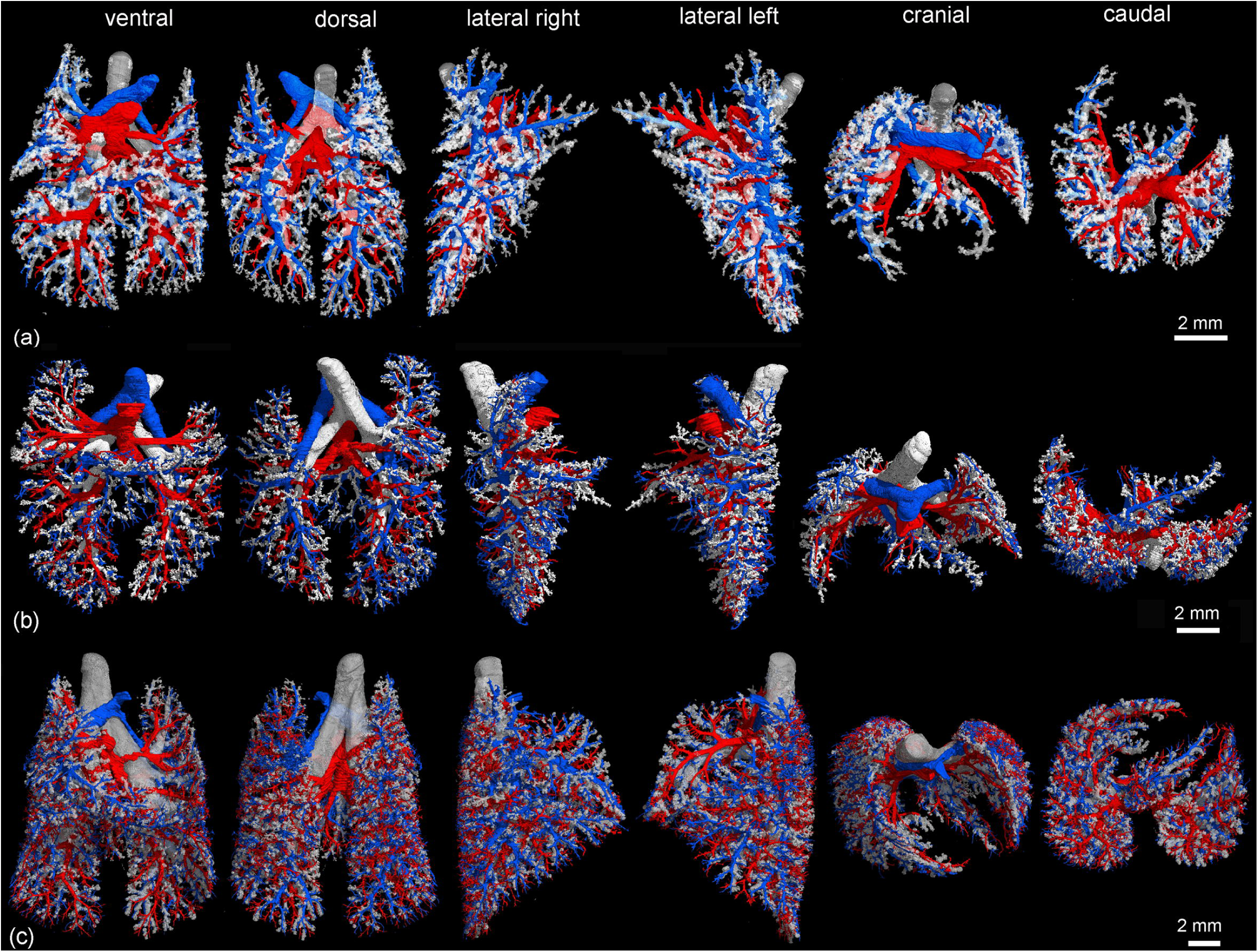
Pulmonary vasculature in relation to the bronchial tree at 35 dpn (a), at 49 dpn (b) and at 57 dpn (c). The lungs are shown from different perspectives: in ventral, dorsal, lateral, cranial and caudal views (from left to right). The pulmonary artery and vein are indicated by blue and red respectively, the bronchial tree appears white.

**Fig. 7.**
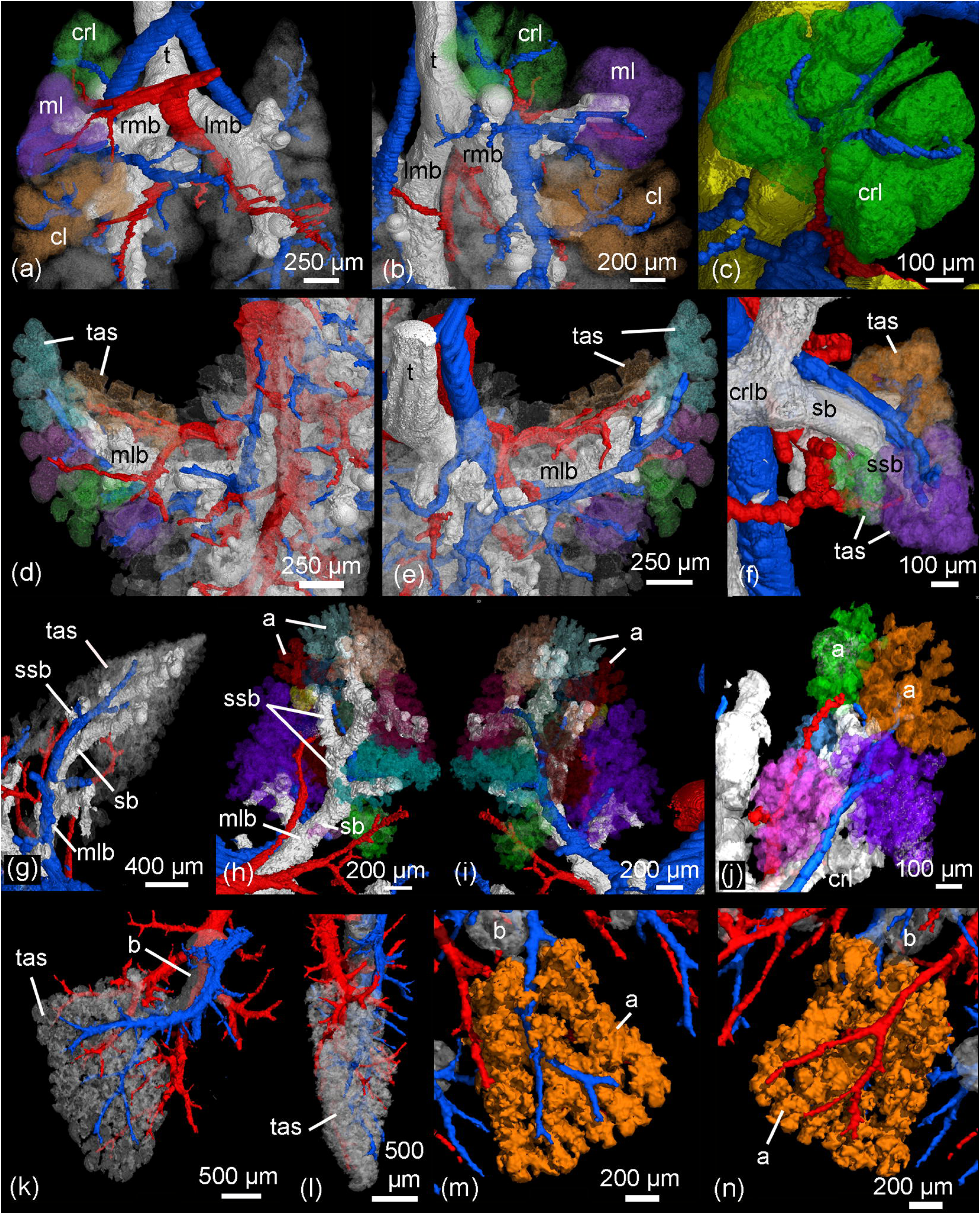
Details of the developing pulmonary vasculature in relation to bronchial tree and terminal air spaces. In the neonate *Monodelphis domestica* (a-c) a simple bronchial tree opens into large terminal air spaces. The pulmonary vasculature exceeds the bronchial tree (a, b) and supplies the terminal air spaces (c). By 4 dpn (d, e) first segmental bronchioles open into several smaller terminal air spaces. The pulmonary artery and vein run in parallel to the bronchial tree and supply the terminal air spaces. By 11 dpn (f) sub-segmental bronchioles supply the saccular terminal air spaces. The pulmonary artery follows the bronchial tree closely and its rami supply the single terminal air spaces. By 21 dpn (g-i) the lung is still at the saccular stage, however the terminal air spaces consist of saccular acini. The pulmonary artery follows the bronchial tree closely, whereas the pulmonary vein reaches the terminal air spaces from the periphery. By 35 dpn (j) the lung is at the alveolar stage and true alveolar acini can be found. The ramification of the pulmonary vasculature increases in order to supply the numerous acini. The highest degree of ramification of the pulmonary vasculature can be seen by 57 dpn (k-n). The pulmonary artery, approaching from the dorsal side, and the pulmonary vein, approaching from the ventral side, form a dense capillary net within the acinus. The pulmonary artery and vein are indicated by blue and red respectively, the bronchial tree is white and the terminal airspaces appear white or colored transparent. Different terminal air spaces/acini are shown with different colors. a, acinus; b, bronchiole; cl, caudal lobe; crl, cranial lobe; crlb, cranial lobe bronchiole; lmb, left main bronchus; ml, middle lobe; mlb, middle lobe bronchiole; rmb, right main bronchus; sb, segmental bronchiole; ssb, sub-segmental bronchiole; t, trachea; tas, terminal air spaces

**Fig. 8.**
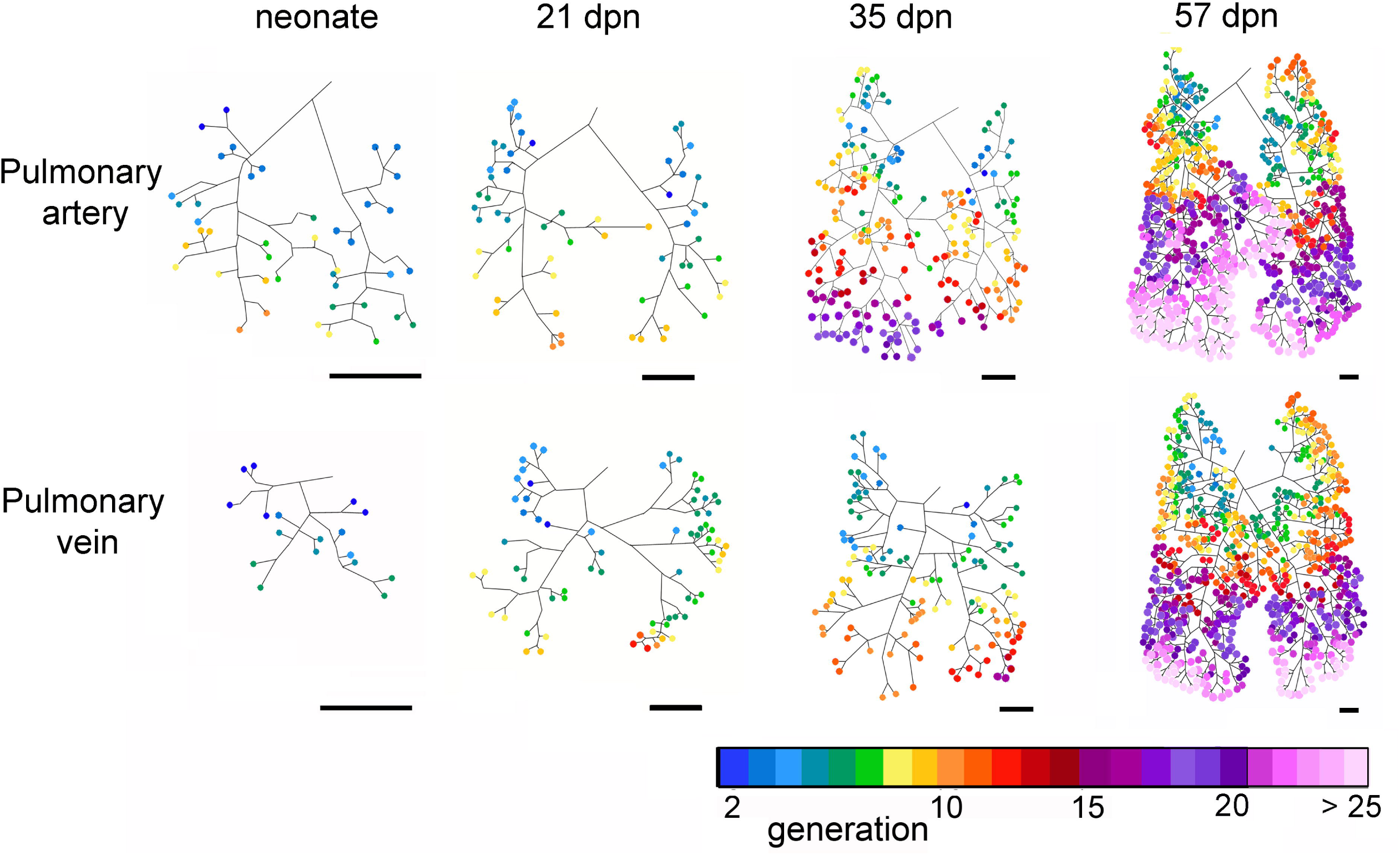
Centerline reconstructions showing the end-generations of the rami of the pulmonary artery and pulmonary vein (rainbow color) in the *Monodelphis domestica* lung in a neonate and by 21 dpn, 35 dpn and 57 dpn. The colored circles indicate generations of each end-ramus when the main trunk of the pulmonary artery and vein was defined as 0^th^ branch. Color bar indicates the end-branching generation with the corresponding color. Scale bar = 1 mm.

**Fig. 9.**
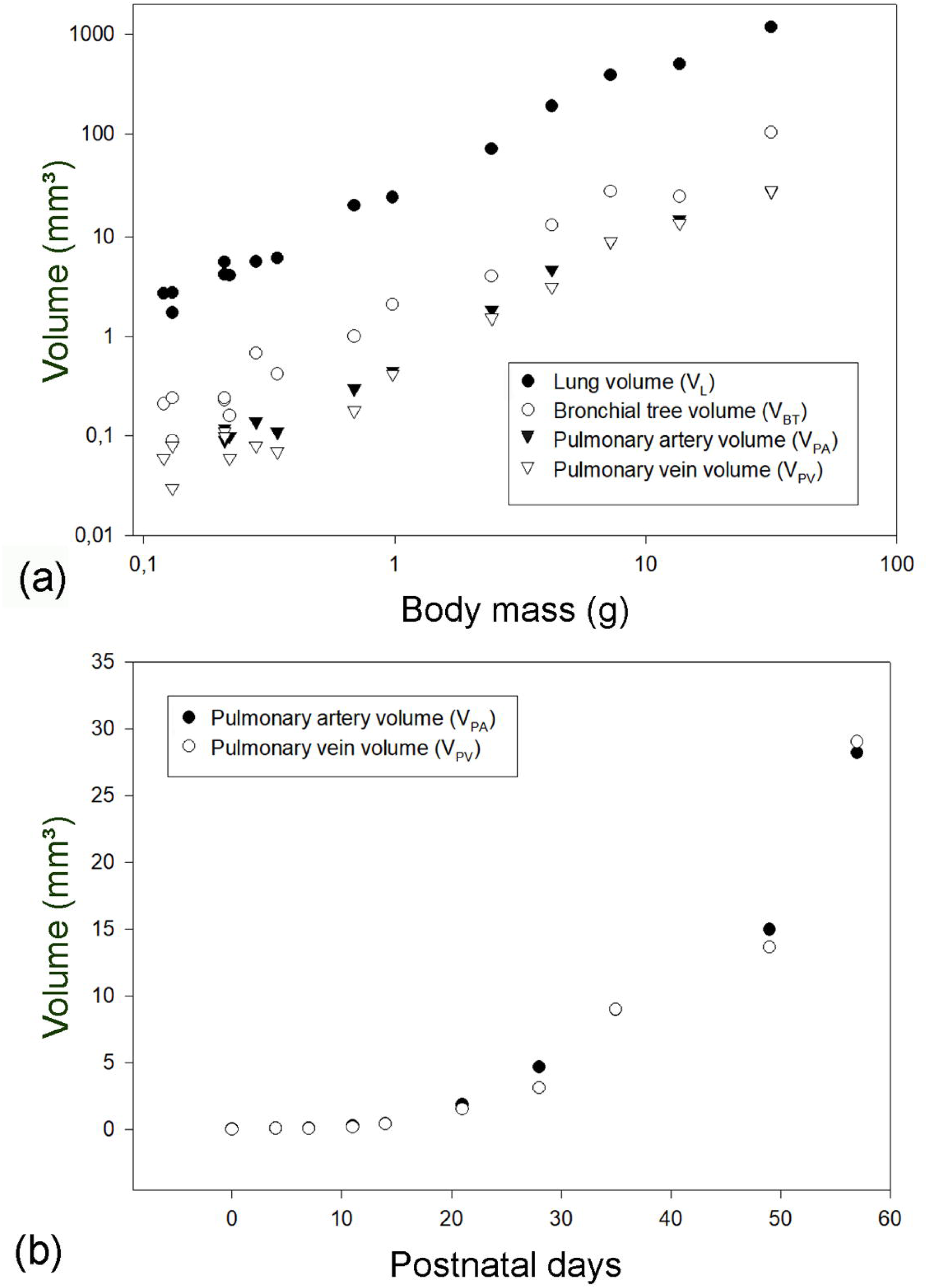
Double logarithmic plots of the lung volume (V_L_), bronchial tree volume (V_BT_), pulmonary artery volume (V_PA_) and pulmonary vein volume (V_PV_) against body mass for *Monodelphis domestica* in the postnatal period (a). Linear plots of the pulmonary artery volume (V_PA_) and pulmonary vein volume (V_PV_) for the postnatal period (b). The graphs are based on individual animal data.

### Development of the pulmonary vasculature

The pulmonary vasculature of the lungs from neonate to 57 dpn showed a marked size increase and a qualitative increase in vascular complexity and density (Fig. 2 a). In the neonate *Monodelphis domestica* the pulmonary vasculature has a simple branching pattern and leads to the periphery of the lung (Fig. 2b). Compared to the bronchial tree, which ends in short lobar bronchioles, the pulmonary vasculature appears to be more developed and extends to the large terminal air spaces (Fig. 4a). The volumes of the pulmonary artery are 0.07 mm³ and 0.06 mm³ respectively (Table 1). Inspection of the lung periphery at higher magnification shows how the pulmonary artery and vein communicate with the large terminal air spaces (Fig. 7 a-c). The centerline reconstruction of the pulmonary vasculature of the neonate lung reveals that the branches of the pulmonary artery have a median generation of 5, a maximum generation of 10 and a total of 41 vessels (Fig. 8, Table 2). The pulmonary vein appears to be less developed than the pulmonary artery and has a mean branch generation of 4, a maximum branch generation of 6 and only a total of 17 vessels.

Between 4 and 7 dpn the pulmonary vasculature exhibits only a small size increase (Fig. 2 c, d; 4 b, c). The pulmonary arteries have a volume of 0.10 and 0.13 mm³ and the pulmonary veins have a volume of 0.09 and 0.08 mm³ (Table 1). The lung reaches the saccular period and the terminal air spaces become more subdivided. The bronchial tree is still dominated by short branching airways opening into terminal saccules. However, compared to earlier developmental stages a more differentiated bronchial tree is present. Segmental bronchioles can be seen by 4 dpn and first sub-segmental bronchioles are evident by 7 dpn. By 7dpn the bronchial tree development has cached up with the pulmonary vasculature. Details of the pulmonary vasculature at the lung periphery by 4 dpn indicate the presence of new generations of branches that communicate with the new formed terminal air spaces (Fig. 7 d, e).

From 11 dpn to 21 dpn, the lung is still at the saccular stage of lung development, however increases markedly in complexity. This increase in complexity is concurrent with an increase in saccular number and decrease in the size of the saccules. The pulmonary vasculature, developing in parallel to the bronchial tree, is characterized by enhanced architectural complexity (Fig. 2 e; 3 a, b; 4 d; 5 a, b). The pulmonary vasculature exhibits a further size increase from 0.30 to 1.68 mm³ for the pulmonary artery and 0.18 to 1.54 mm³ for the pulmonary vein. Details of the lung periphery reveal that by 11 dpn sub-segmental bronchioles supply the saccular terminal air spaces. The pulmonary artery follows the bronchial tree closely and its rami supply the single terminal air spaces (Fig. 7 f). By 21 dpn the lung is still at the saccular stage, however the terminal air spaces consist of saccular acini. The pulmonary artery follows the bronchial tree closely, whereas the pulmonary vein reaches the terminal air spaces from the periphery (Fig. 7g-i). The structural complexity of the pulmonary vasculature increases moderately (Fig. 8). By 21 dpn the pulmonary artery and pulmonary vein have a median branch generation of 6, a maximum branch generation of 10 and 12, and a total number of vessels of 56 and 70 respectively (Table 2).

Between 28 and 35 dpn, substantial changes take place in the lung of *Monodelphis domestica*. With the start of alveolarization at 28 dpn, the lung is at the transition from the saccular to the alveolar stage of lung development and by 35 dpn the lung has fully reached the alveolar stage of lung development. During this period the pulmonary vasculature exhibits a marked volume increase from 4.68 to 8.97 mm³ for the pulmonary artery and 3.15 to 9.03 mm³ for the pulmonary vein (Table 1). The complexity of the pulmonary vasculature and of the bronchial tree is increasing furthermore (Fig. 3 c, d; 5 c; 6 d). By 35 dpn the well-developed bronchial tree extends to the new formed alveolar acini. The ramification of the pulmonary vasculature increases in order to supply the numerous acini (Fig. 7 j). The centerline reconstruction of the pulmonary vasculature at 35 dpn reveals that the vessel density increases. The branches of the pulmonary artery and pulmonary vein have a median generation of 10 and 8, a maximum generation of 20 and 14 and a total number of 202 and 118 vessels (Fig. 8, Table 2).

The pulmonary vasculature of the lung at 49 and 57 dpn shows a marked increase in vascular density (Fig. 3 e, f). The mature bronchial tree, accompanied by the pulmonary vasculature, is still extending by forming new bronchioles, meeting the requirements of the growing lung (Fig. 6 b, c). Details of the pulmonary vasculature of the terminal air spaces at the lung periphery by 57 dpn reveal a high degree of ramification of the pulmonary vasculature (Fig. 7 k-n). The pulmonary artery, approaching from the dorsal side, and the pulmonary vein, approaching from the ventral side, form a dense capillary net within the acinus. The branching patterns of the pulmonary artery and of the pulmonary vein fill the entire lung and appear highly organized. The centerline reconstruction of the pulmonary vasculature at 57 dpn shows the highest vessel density. The branches of the pulmonary artery and pulmonary vein have a median generation of 14 and 12, a maximum generation of 27 and 25 and a total number of 695 and 644 vessels (Fig. 8, Table 2).

## Discussion

Viability of the newborn mammal depends on an adequately developed respiratory apparatus of the neonate (Frappell & Mac Farlane, 2006). Although the lungs of newborn marsupials are structurally immature, in the majority of marsupial species, the lungs become ventilated with birth and take over respiratory function. Depending on the developmental degree of the lung, pulmonary respiration is supported by cutaneous respiration to a higher or lower amount (Mortola et al., 1999; MacFarlane & Frappell, 2001; Mac Farlane et al., 2002; Frappell & Mac Farlane, 2006; Simpson et al., 2011; Ferner, 2018, 2021b).

The lung in the newborn gray short-tailed opossum has large terminal air spaces. The cranial, middle and accessory lobes of the right lung consist of one large terminal air space respectively (Ferner, 2024). Although the air space septa consist of capillaries on both sides, forming a blood-air-barrier facilitating gas exchange, a continuous double capillary septum is not present yet, attributing the lung of the newborn gray short-tailed opossum to the late canalicular stage (Modepalli et al., 2018, Ferner, 2024).

Even if the gray short-tailed opossum represents a basal mammalian species, the pulmonary vasculature bears some similarities to the pulmonary vasculature of the mouse, rat and human lung (Frey et al., 2004; Netter & Mühlbauer, 2011; Razavi et al., 2012; Phillips et al., 2017).

Most of our knowledge about structural changes in lung development stem from eutherians, especially rat models (Burri, 1974; Kauffman et al., 1974), and humans (Weibel, 1967; Hislop & Reid, 1972). However, some studies investigated marsupial species for postnatal lung development (Runciman et al., 1998a; Burri et al., 2003; Ferner, 2021a). These studies mainly used histology and morphometric methods, that are based on a random or stereological sampling of a planar section of the lung vasculature to make conclusions about the lung as a whole. Due to the complexity of the pulmonary circulation with many generations of branches and a large number of vessels it has proven challenging to evaluate the whole pulmonary vasculature of the lung. In the morphometric studies of marsupial lung development there is no differentiation between pulmonary artery and pulmonary vein and usually only volume fractions and volumes for blood vessels have been reported. In the quokka, the non-parenchymal vascular volume increase accelerated in the successive developmental stages while the airway and connective tissue volumes progressed in a decreasing order from canalicular to alveolar stage (Burri et al., 2003). Runciman et al. (1998a) described a significant increase in the large blood vessel fraction of the non-parenchyma in the 180-day old pouch young and adult Tammar wallaby. This is similar to findings in the developing rat lung (Burri et al., 1974). It can be assumed that, as the lung changes from the saccular to the alveolar stage, the emphasis shifts from conducting airway development to development of the pulmonary vasculature (Runciman et al., 1998a).

The pulmonary vasculature in the lung of *Monodelphis domestica* follows the course of the bronchial tree and corresponds to the six pulmonary lobes. The lobation of the lung is similar to previous descriptions of the monotreme short-beaked echidna (Perry et al., 2000) and many marsupial and eutherian species (Nakakuki, 1980; Cope et al., 2001, Ferner & Mahlow, 2023). The asymmetry of the right (four lobes) and the left lung (two lobes), as present in *Monodelphis domestica*, is common in other basal taxa and many distantly related mammalian groups and can be considered as plesiomorphic for Mammalia (Ferner & Mahlow, 2023).

The anatomy of the pulmonary artery and pulmonary vein present in the lung of *Monodelphis domestica* resembles the pulmonary vasculature described in another marsupial species, the Virginia opossum (*Didelphis virginiana*) (Cope et al., 2001). A comparison with reconstructions of the pulmonary vasculature using corrosion casts and µCT in the pig (Nakakuki, 1994a), cow (Nakakuki, 1994b), Japanese deer (Nakakuki, 1993), rabbit (Autifi et al., 2015), mouse and rat (Counter et al., 2013), reveals overall similar trees for the pulmonary artery and pulmonary vein. Some variations in the pulmonary vasculature have been described for the Japanese monkey (Nakakuki, 1986) and the orangutan (Nakakuki & Ehara, 1991).

This is the first study providing three-dimensional reconstructions of the pulmonary vasculature of the developing lung in a marsupial species. Information about the structural modifications of the cardiovascular circulation in newborn marsupials are sparse. At birth, the ductus arteriosus and interventricular communication close within hours of birth, though an interatrial communication via a fenestrated septum persists for 2-3 days (Gannon et al., 1989; Runciman et al., 1995; Ferner, 2021b). The persistence of the right-to-left shunt presumably prevents full expansion of the pulmonary blood vessels. This could explain why the volume density of blood vessels in the tammar wallaby was about an eight of the volume density of the airways in the non-parenchymal portion of the lung in the newborn tammar (Runciman et al., 1998b). In contrast, the volume densities of the airways and blood vessels were approximately equal in the newborn rat (Frappell & MacFarlane, 2006). The unique situation of the persistence of a right-to-left shunt prevents discreet pulmonary and systemic circuits, permits blood to bypass the lungs and allows venous admixture of blood low in oxygen content. This might be necessary to allow for cutaneous respiration during the first postnatal days in marsupials (Ferner, 2021b).

The pulmonary vasculature in the gray short-tailed opossum goes through a remarkable structural development from poorly branched trees in the neonate lung to a highly ramified pulmonary artery and pulmonary vein that fill the entire lung by 57 dpn. Comparable studies of the developing pulmonary vasculature in marsupials are missing so far. However, a few studies using µCT investigated the postnatal development of the pulmonary vasculature, in particular the pulmonary artery, in eutherian species (Razavi et al., 2012; Phillips et al., 2017). Similarly to the gray short-tailed possum, reconstructions of the mouse pulmonary vasculature at 2 and 4 weeks, and at 3 months of age showed a marked increase in vascular complexity and density (Phillips et al., 2017) with age. The largest increase in blood vessel number was observed between 2 weeks and 4 weeks and between the 21^st^ and 40^th^ generation, with vessels having diameters of 75µm and below (Phillips et al., 2017). Also, in the gray short-tailed opossum the vessel number in the vascular trees of the pulmonary artery and pulmonary vein increased markedly during the postnatal period, in particular by 35 dpn, when alveolarization started and by 57 dpn, when the final adult lung structure was attained.

This study did not estimate diameters or lengths of the blood vessels. However, other studies reported no differences in vessel length with age or according to diameter (Counter et al., 2013; Phillips et al., 2017). Branching in the pulmonary vasculature seems to be fractal in nature and the length of vessel segments between branch points is consistent throughout the vascular tree (Frey et al., 2004). Therefore, complexity in the pulmonary vasculature is obviously not determined by vessel length, but by the degree of progressively increasing vascular branching (Weibel et al., 1963).

This study has limitations. Since the cardiovascular system was not perfused, the blood vessels were partly filled with blood. This caused difficulties during the segmentation process, since different gray values resulted from air-filled and blood-filled vessels. When a change in the gray values occurred, the region grower was corrected for the new value. In addition, there were differences in the resolution of the scanned lungs, affecting the segmentation of the pulmonary vasculature. In all lungs the pulmonary vein and pulmonary artery was segmented as far as possible to the periphery of the lung. A final limitation, µCT scanners are expensive. But if available, high-resolution µCT imaging and 3D reconstruction offer new possibilities to characterize the structural changes associated with lung development.

## Conclusion

In the marsupial gray short-tailed opossum the process of vascular genesis, which takes place intrauterine in the placental fetus, is shifted to the postnatal period and therefore more easily accessible for investigation. The development of the pulmonary vasculature from a simple vascular tree consisting of only a few generations of vessels to the final densely vascularized tree characterized by high branching complexity, takes place during the functional state in a continuous morphogenetic process. Similarities to the vascular genesis described in other marsupials and eutherians suggest that this process is highly conservative within mammalian evolution. Lung development, including vascular genesis, in placental and marsupial mammals follows similar patterns.

## Author contributions

**KF:** design, dissection of lungs, 3D-reconstructions, data collection and analysis/interpretation, manuscript writing and editing

## Acknowledgements

I am very grateful to Kristin Mahlow (lab manager of CT-lab) for staining, sample preparation and acquisition of µCT-scans. I thank the animal keepers Petra Grimm and Annett Billepp as well as Dr. Peter Giere and Dr. Peter Bartsch of the animal facility of the Museum für Naturkunde Berlin for breeding and providing the animals for this project. I acknowledge the financial support from the German Research Foundation (DFG) with the module “temporary position for the principal investigator” (Grant No. FE1878/2-1). Open access funding enabled and organized by Projekt DEAL.

## Data availability statement

The data that support the findings of this study and additional images and videos of 3D reconstructions of the pulmonary vasculature are made publicly available with figshare (data: https://doi.org/10.6084/m9.figshare.25610403; original images: https://doi.org/10.6084/m9.figshare.25610466; 3D-images: https://doi.org/10.6084/m9.figshare.25610493 and 3D-videos: https://doi.org/10.6084/m9.figshare.25610790).

